# Folic acid prevention of neural tube defects requires retinoic acid produced by ALDH1L1

**DOI:** 10.1101/2025.10.14.681787

**Authors:** Tamir Edri, Tali Abbou-Levy, Dor Cohen, José M. Inácio, Yehuda Shabtai, Graciela Pillemer, José António Belo, Abraham Fainsod

## Abstract

Folic acid (FA) supplementation during pregnancy is the commonly accepted treatment to prevent neurodevelopmental defects. The mechanism by which FA prevents neural tube defects (NTDs) remains unclear. FA also prevents other developmental malformations, including the alcohol-induced malformations in Fetal Alcohol Syndrome models. We show that FA acts through a metabolic link to retinoic acid (RA) signaling. Using a *pax3*-knockdown *Xenopus* model of FA-rescueable NTDs, we show that RA or its precursors equally rescue these defects. Similarly, FA rescues alcohol-induced NTDs in a model previously shown to be rescued by RA. We identify the FA-metabolizing enzyme, formyl tetrahydrofolate dehydrogenase (ALDH1L1, FTHFD), encoded by the *aldh1l1* gene, as essential for this rescue. Mechanistically, FA upregulates *aldh1l1* expression, leading to increased RA biosynthesis. Knockdown of the ALDH1L1 activity using CRISPR/Cas9 abolishes the FA protective effect. To support these observations, we show that the human ALDH1L1 enzyme converts retinaldehyde to RA, and its overexpression restores neural tube closure in *aldh1l1*-knockdown embryos when retinaldehyde is provided. At the cellular level, reduced RA signaling induces an overproliferation of neural plate precursors and a pathological expansion of the neural tube. ALDH1L1 enables FA to restore normal neural plate proliferation, thereby preventing NTDs. These findings establish ALDH1L1 as a previously unrecognized enzymatic link between FA (vitamin B9) and RA signaling, revealing how FA supplementation safeguards neural development and suggesting opportunities to refine strategies for NTD prevention.

**Significance Statement:** Despite the global success of folic acid (FA) supplementation in preventing neural tube defects (NTDs), the medical community continues to debate its exact mechanism, optimal dosage, and why it fails in some cases. This study provides a breakthrough by providing a mechanistic explanation linking FA supplementation and retinoic acid (RA) signaling. We identify the enzyme ALDH1L1 as the molecular bridge between FA and RA and demonstrate that FA protection is indirect, requiring ALDH1L1 to convert Vitamin A into RA. This discovery reframes the debate surrounding FA in the prevention of NTDs. Clinically, our findings suggest that integrating FA supplementation with optimized Vitamin A levels could improve current preventive practices and maximize neurodevelopmental safeguards during early pregnancy.

## INTRODUCTION

Approximately 6% of births worldwide exhibit developmental defects (1), among them, debilitating neural tube malformations (2). Failure of neural tube closure (NTC) during early embryogenesis results in neural tube defects (NTDs) that can occur at any level of the central nervous system (CNS), with an incidence of 19 per 10,000 newborns (1, 2). NTD induction has been linked to abnormal folic acid (FA, vitamin B9) metabolism, impaired neural tube morphogenesis, cell migration, Wnt/Planar Cell Polarity (PCP) signaling, retinoic acid (RA) signaling, and epigenetics (1–4), but the teratogenic mechanism remains unclear. NTD risk is influenced by genetic and environmental factors, such as medications and toxins (3, 5). Prenatal alcohol exposure, especially Fetal Alcohol Syndrome (FAS), the severe form of Fetal Alcohol Spectrum Disorder (FASD), increases NTD risk (6–9). In 1991, the MRC Vitamin Study Research Group concluded that FA supplementation reduces NTD incidence, a finding supported by others (10). FA lowers NTD risk but doesn’t eliminate it, suggesting multiple mechanisms (3, 4). After food fortification was implemented, concerns about toxicity and teratogenicity from high FA levels were raised (11, 12). Understanding how FA decreases NTDs is vital to understanding the role and potential risks of FA supplementation.

Lowering RA levels leads to FAS-like malformations, and supplementing EtOH-induced malformations with vitamin A (retinol, ROL) or retinaldehyde (RAL), the RA precursors, rescues them, showing that FAS arises from diminished retinoic acid (RA) signaling during embryogenesis (13). Acetaldehyde, from ethanol (EtOH) oxidation, competes with RAL, the oxidation product of ROL, for the embryonic retinaldehyde dehydrogenase 2 (RALDH2, ALDH1A2) activity (14–17). This competition reduces embryonic RA production (18), leading to teratogenic effects indistinguishable from those of FAS (13). We previously showed that reducing RA biosynthesis with EtOH or RALDH inhibitors causes NTC defects in *Xenopus* embryos, which can be rescued with ROL or RAL (8). Now, we show that FA also rescues these defects. Similar rescue results were observed in a pax3 knockdown model (CRISPR/Cas9). The developmental rescue window of FA overlaps the retinoid rescue window. We analyze the *aldh1l1* gene, which encodes 10-formyltetrahydrofolate dehydrogenase (Aldh1l1, Fthfd), with a conserved carboxy-terminal domain similar to that of retinaldehyde dehydrogenases (19, 20) that oxidize RAL to RA (19, 20). Using CRISPR/Cas9, we show that Aldh1l1 activity is essential for FA rescue of NTC defects caused by inhibition of RA biosynthesis or *pax3* targeting. Human ALDH1L1 overexpression rescues *aldh1l1* knockdowns when supplemented with RAL. We also show, using enzymatic kinetics, that human ALDH1L1 produces RA *in vitro*. FA exposure induces ectopic *aldh1l1* expression. This molecular pathway is conserved in mouse cells, including *Aldh1l1* upregulation and RA production. These results suggest *ALDH1L1* is crucial for FA-mediated NTD rescue involving RA biosynthesis.

## RESULTS

### FA rescues NTC defects in reduced RA and pax3 knockdown models

To study NTD rescue by FA during pregnancy, we established two experimental models to induce NTC defects: *pax3* knockdown or inhibition of RA biosynthesis. In the first model, we targeted the *pax3* gene (CRISPR/Cas9) in *Xenopus* embryos to create an *Xsplotch* model equivalent to the *splotch* mouse, which is rescued by FA (21). To generate *pax3* CRISPant embryos, we chose an exon 1 sgRNA that induces indels in both L and S *pax3* homoeologs (Supplementary Fig. 1A). TIDE software deconvolution analysis (22) revealed indel efficiencies of over 82% and frameshift efficiencies of 63.1% and 73.2% for the L and S genes, respectively (Supplementary Fig. 1B-E). CRISPant embryos showed a significant reduction in *pax3*.L and *pax3*.S expression from late gastrula (NF12) to NTC stages (NF19) as determined by qPCR (Supplementary Fig. 1F, G). *Pax3* CRISPant embryos exhibited NTC defects during post-closure stages (NF19-20) (8) when stained with *sox3* (Fig. 1A, A’, B, B’) or *shh* (Fig. 1C, C’, D, D’). We scored for open neural tubes at closure stages to quantify the induction of NTC defects. *Xsplotch* embryos exhibited a significantly higher incidence of NTC defects (Fig. 1E). These findings suggest that *pax3* CRISPants develop NTC defects similar to mouse *splotch* mutants.

**Figure 1.**
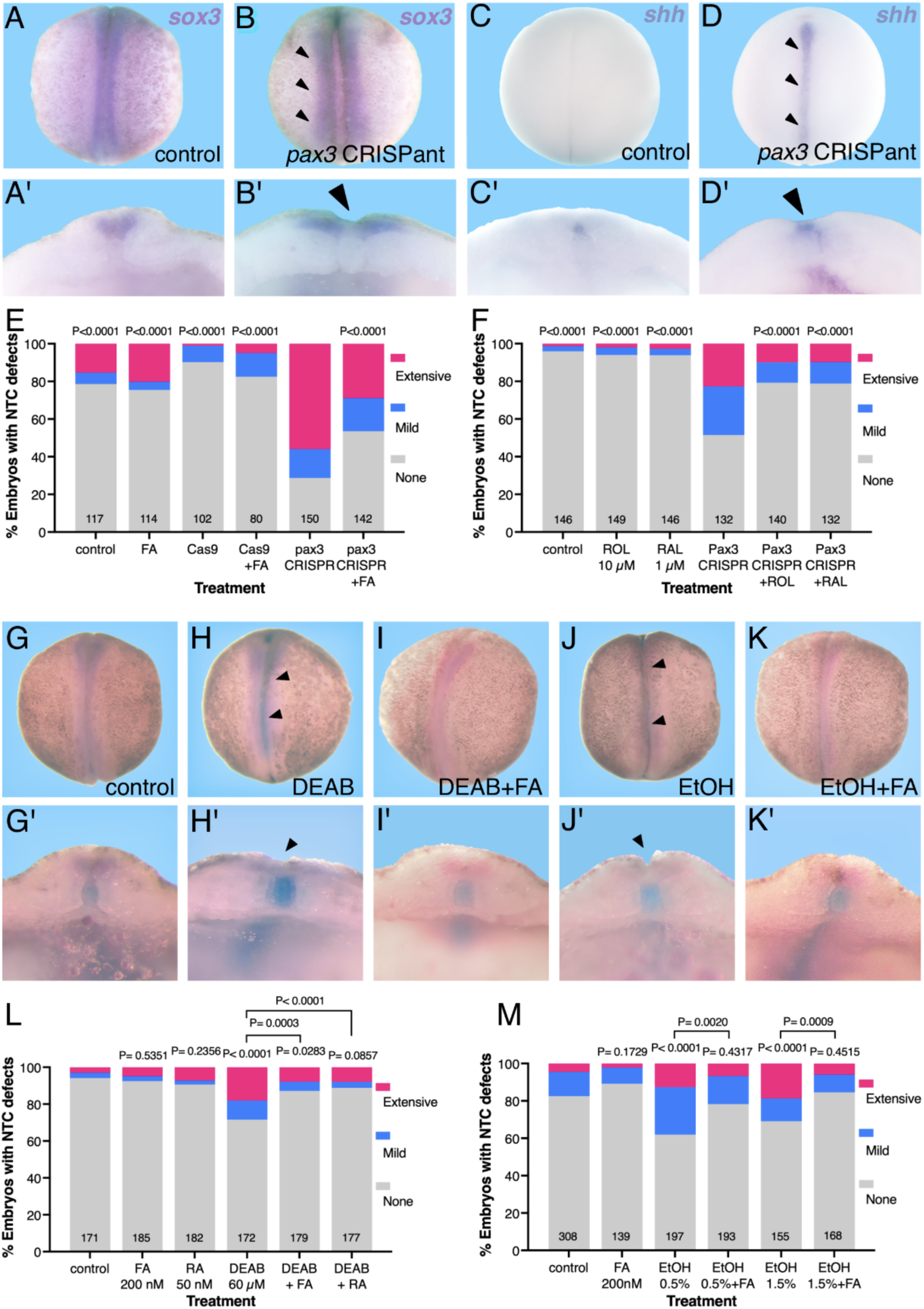
Neural tube closure defects in *pax3* CRISPants and reduced RA embryos are rescued by FA and RA precursors. *Xsplotch* and embryos with reduced RA signaling (DEAB 60 µM or EtOH 0.5% and 1.5% vol/vol) (8), were analyzed for NTC defects and their rescue by FA (200 nM), RA (50 nM), ROL (10 µM), or RAL (1 µM) supplementation. (A-D’) Embryos CRISPant for *pax3* exhibit neural tube closure defects (arrowheads). Analysis by whole-mount *in situ* hybridization using *sox3* to mark the neural plate (A-B’) or *shh* to label the notochord and floor plate (C-D’) probes to visualize the open neural tubes. (A-D) Dorsal views of embryos at NTC closure stages (NF19/20). (A’-D’) Cross-section of the same embryos bisected to demonstrate the open neural tube (B’, D’). Quantitation of the FA (E) and ROL and RAL (RA precursors)(F) rescue of NTC defects in *pax3* CRISPants. (G-K) Embryos treated to reduce RA signaling (DEAB or EtOH) were analyzed at NF19/20 for the incidence of NTC defects (arrowheads) by whole-mount *in situ* hybridization, using pax3 (magenta) to label the neural plate border and *chrd* (dark blue) to mark the notochord, which is visible when the neural tube remains open. (G’-K’) Cross-sections of the same embryos shown in G-K. (L) Quantitative analysis of NTC defect induction by DEAB and FA rescue. (M) NTC defect induction and rescue in embryos treated with EtOH (0.5% or 1.5%) and FA. The NTC defect incidence is shown as the percent of affected embryos. Sample sizes are shown, and statistical significance was calculated using Fisher’s exact test relative to either the *pax3* CRISPant or the control sample in the reduced RA (DEAB or EtOH) experiments.

To rule out a developmental delay in *pax3* CRISPants as the source of open neural tubes, we analyzed embryos from independent batches (N=5) at early neurula (NF13) and re-analyzed the same embryos at neural tube closure (NF19 and NF21) stages. When control embryos reached late gastrula (NF12) or early neurula (NF13), many *pax3* CRISPant embryos were still at mid-gastrula (NF10.5-NF11.5), indicating a developmental delay during gastrulation (Supplementary Fig. 2A). Re-analysis of the same embryos at NF19 and NF21 showed no difference in the stage distribution of the *pax3* CRISPants compared to their control siblings (Supplementary Figs. 2B, C). These findings support a recovery of the *pax3* CRISPants and the closing of the developmental delay.

To study the FA rescue of NTC defects in *Xsplotch* embryos, CRISPants were treated with FA and analyzed for neural tube closure. Several FA concentrations were tested for efficient rescue of *pax3* CRISPant NTDs (Supplementary Fig. 3A). FA supplementation (200 nM) significantly reduced NTC defects in *pax3* CRISPant embryos (Fig. 1E). Controls with FA only and Cas9 injections showed no significant NTC defect induction (Fig. 1E). These results validate the *pax3* CRISPant experimental model for FA rescue of NTC defects in *Xenopus*, similar to mouse *splotch*.

In the second experimental model, we induced NTC defects by pharmacologically inhibiting RA biosynthesis (8), thereby allowing us to elucidate the molecular biochemical mechanism of NTD rescue by FA. *Xenopus* embryos were treated with the RALDH inhibitors, DEAB (4-diethylaminobenzaldehyde) (23) or EtOH (0.5% or 1.5% vol/vol) (17), from midblastula (NF8.5). NTC defects were analyzed at closure stages (NF19-20) by staining the neural plate with *pax3* or the notochord with *chrd* (Fig. 1G-K’). Both RA biosynthesis inhibitors (DEAB or EtOH) induced NTC defects efficiently (Fig. 1G, G’, H, H’, J, J’, L, M). Focusing on reduced RA induction of NTC defects, after titration, a FA concentration (200 nM) that consistently up-regulates RA-target genes was selected (Supplementary Fig. 3B-D). Supplementing with FA efficiently prevented NTC defects in embryos with reduced RA (Fig. 1I, I’, K, K’, L, M). As a control, DEAB-treated embryos were also supplemented with RA (50 nM), which rescued NTC defects by compensating for decreased RA biosynthesis (Fig. 1L). The FA rescue of NTDs induced by reduced RA biosynthesis suggests a biochemical mechanism in which FA promotes RA production.

### Rescue of NTC Defects by Folic Acid Involves Retinoic Acid Biosynthesis

To further support the connection between FA supplementation and RA signaling, we rescued *Xsplotch* embryos with the RA precursors, ROL or RAL (Fig. 1F). Adding ROL (10 µM) or RAL (1 µM) efficiently reduced NTC defects in *pax3* CRISPants. Control embryos treated with ROL or RAL didn’t show a significant increase in NTC defects compared to untreated control embryos (Fig. 1F). These findings show that retinoid supplementation reduces NTC defects in *pax3* CRISPants, similar to the mouse *splotch* model (24).

NTC defects induced by reduced RA signaling are most efficiently rescued by providing RA or its precursors during early gastrulation (8). Therefore, we determined whether the FA and retinoid rescues share the same developmental window for rescuing NTDs in *pax3* CRISPants. Adding FA or RAL to *Xsplotch* embryos during midblastula (NF8.5) significantly reduced NTC defects compared to untreated CRISPant siblings (Fig. 2A). Delaying FA or RAL supplementation to NF10.25 diminished rescue efficiency, and treatments starting from mid-gastrula (NF11) had no effect (Fig. 2A). Similarly, FA rescue of EtOH-treated (0.5% or 1.5%) embryos was significant when added during midblastula (NF8.5) stages (Fig. 2B, C), but decreased during early gastrula (NF10.25) or mid-gastrula (NF11) stages (Fig. 2B, C). These findings demonstrate that FA has the same developmental rescue window as RA or retinoids for NTC rescue, supporting a common biochemical mechanism.

**Figure 2.**
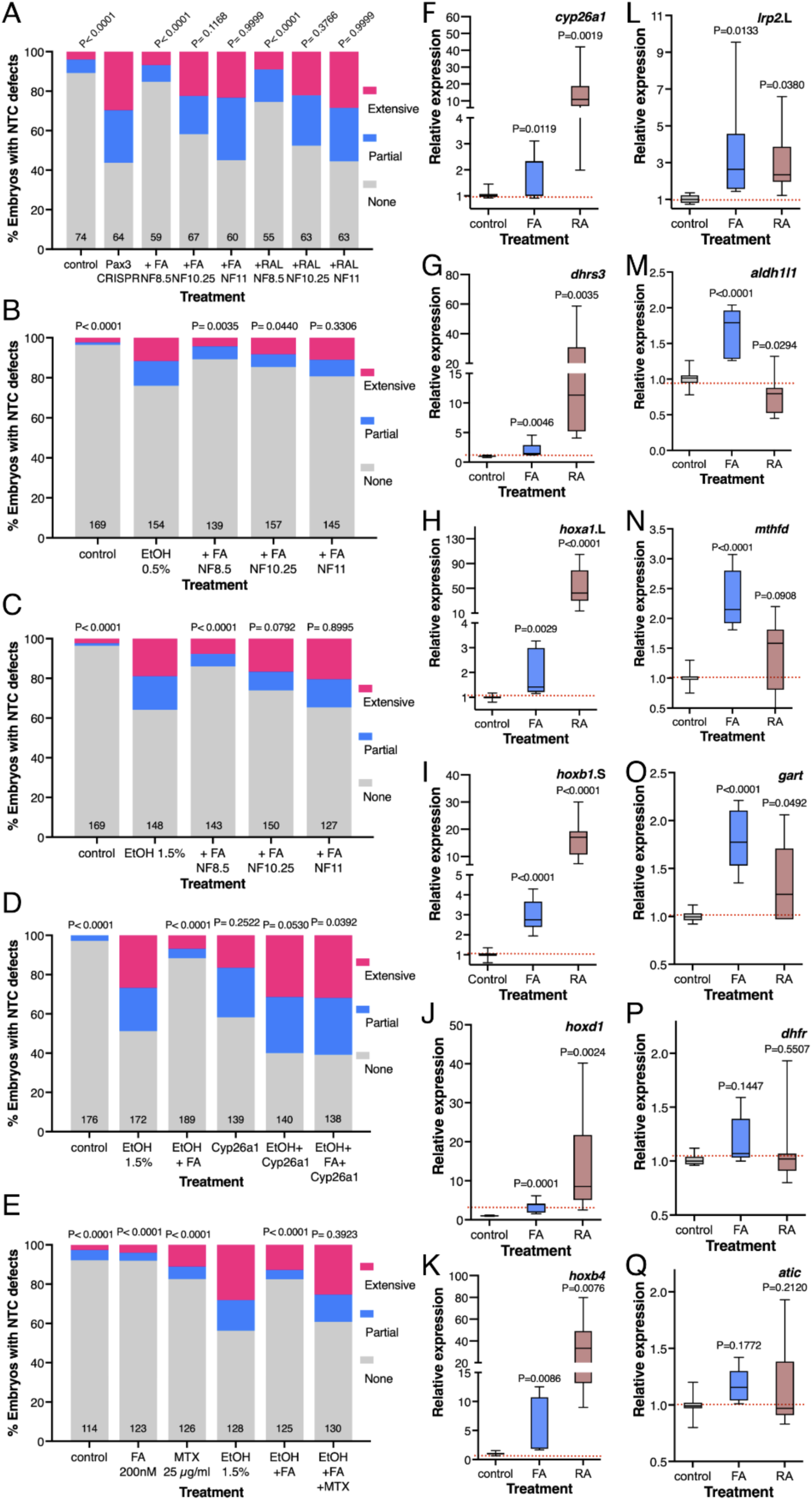
FA supplementation rescues NTC defects, self-regulates its metabolism, and upregulates RA target genes during early gastrulation. (A-C) The developmental window for rescuing NTC defects was studied by administering FA or RAL at various developmental stages: late blastula (NF8.5), early gastrula (NF10.25), and mid-gastrula (NF11). The NTC defect incidence was calculated as the percentage of embryos with partial or extensive NTC defects. Sample sizes are shown, and statistical significance was determined using Fisher’s exact test. (A) Developmental time window analysis of *pax3* CRISPants with FA or RAL. (B, C) FA rescue of embryos treated with 0.5% or 1.5% EtOH. (D) EtOH-treated embryos overexpressing the RA-hydroxylase, Cyp26a1, were treated with FA to determine the involvement of RA in the FA rescue. The NTC defect incidence was calculated as the percentage of embryos with partial or extensive NTC defects. Sample sizes are shown, and statistical significance was determined using Fisher’s exact test. (E) To investigate whether FA must be converted to THF, embryos were also treated with MTX (25 µg/ml) to inhibit DHFR activity, and the percentage of embryos with NTC defects was calculated. (F-Q) Embryos were treated with FA (200 nM) or RA (50 nM) and allowed to develop to early or mid-gastrula (NF10.5). (F-K) Changes in expression levels of RA target genes (*hoxa1*, *hoxb1*, *hoxd1*, *hoxb4*, *cyp26a1*, *dhrs3)* were determined by qPCR. (L-Q) The self-regulatory effect of FA on genes involved in FA metabolism (*lrp2*, *dhfr*, *mthfd*, *gart*, *aldh1l1*, *atic*) was investigated by qPCR. Box-and-whisker plots (5-95 percentiles). Statistical significance was calculated using one-way ANOVA compared to the control expression.

The FA rescue of NTC defects induced by decreased RA biosynthesis (Fig. 1L, M) suggests that FA enhances RA production. To support this observation, we overexpressed the RA hydroxylase Cyp26a1 to metabolize the RA produced (25) and to rule out alternative mechanisms. For biochemical clarity, NTC defects were induced in embryos with EtOH (1.5%), and Cyp26a1 overexpression was followed by FA treatment. Cyp26a1 overexpression completely abolished the FA-rescuing effect compared to FA alone (Fig. 2D), supporting the role of RA in FA rescue. We also analyzed RA target genes in embryos treated with FA or RA. Analysis (qPCR) of RA-responsive *Hox* genes (*hoxa1*, *hoxb1*, *hoxd1*, *hoxb4*) and RA metabolism genes (*cyp26a1*, *dhrs3*) at NF10.5 revealed significant upregulation of all these genes following FA exposure, and RA treatment (Figs. 2F-K and Supplementary Fig. 3B-D). These results support an increase in RA levels following FA treatment, consistent with FA-driven RA biosynthesis rescuing NTC defects.

### ALDH1L1 Is Essential for the Rescue of NTC Defects by Folic Acid

To elucidate the mechanism of FA-promoted RA production, we investigated the potential involvement of enzymes in one-carbon metabolism. FA is converted to tetrahydrofolate (THF) by dihydrofolate reductase (DHFR), an enzyme inhibited by methotrexate (MTX) (25). Adding MTX (25 µg/ml) to embryos treated with EtOH (1.5%) to induce NTC defects significantly blocked the rescuing effect of FA (Fig. 2E), showing that FA must be oxidized to THF to rescue NTC defects. To investigate the self-regulatory effect of FA on enzymes involved in FA metabolism, we analyzed gene expression during gastrula stages in embryos treated with FA or RA. FA treatment upregulated the *mthfd*, *gart*, and *aldh1l1* genes (Figs. 2M-O and Supplementary Fig. 3D). FA also upregulated *lrp2*, which participates in folate uptake and NTD induction (Fig. 2L) (26). However, FA did not significantly upregulate the *dhfr* or *atic* genes (Figs. 2P, Q). RA only significantly upregulated *lrp2* and *gart*, while *aldh1l1* was downregulated by RA (Figs. 2L, M, O, and Supplementary Fig. 3E). These results show that FA supplementation triggers a complex self-regulatory response in the FA metabolic network during early gastrulation.

The folate-metabolism enzyme, 10-formyltetrahydrofolate dehydrogenase (Aldh1l1, Fthfd), encoded by the *aldh1l1* gene, exhibits extensive sequence homology (∼48%) with NAD-dependent aldehyde dehydrogenases (ALDH) that produce RA (19). This extensive homology supports the hypothesis that Aldh1l1 could mediate the FA-promoted RA production. In agreement, during gastrulation, the *aldh1l1* gene is up-regulated by FA (Fig. 2M and Supplementary Fig. 3E). To study Aldh1l1 as the enzyme mediating the FA rescue effect, we targeted exon 14 of the *aldh1l1* gene in *Xenopus* embryos using CRISPR/Cas9 (Supplementary Fig. 4A). DNA from neurula-stage CRISPants was sequenced to validate the disruption of *aldh1l1* (Supplementary Fig. 4B, C). TIDE decomposition analysis (22) revealed an indel efficiency above 88% and a frameshift efficiency of 65.6% (Supplementary Fig. 4D). The CRISPant embryos exhibited a significant reduction in *aldh1l1* expression during late gastrula (NF12) and early neurula (NF14-15; Supplementary Fig. 4E).

Double CRISPant embryos targeting both *aldh1l1* and *pax3* were generated to study the mechanism of FA rescue. The NTC defects in *pax3* CRISPants were efficiently rescued by FA treatment (Figs. 1E, 3A-C’). In double, *aldh1l1* and *pax3* CRISPants, FA failed to rescue the NTC defects (Fig. 3D, D’, E, E’). These findings support that a fully functional *aldh1l1* gene is essential for FA to prevent NTC defects in *Xsplotch* embryos. Quantifying the *aldh1l1* requirement showed that the loss of Pax3 activity induced NTC defects, and FA alleviated this effect (Fig. 3F). While double *aldh1l1*+*pax3* CRISPants still showed a high NTC defect incidence, which FA supplementation did not improve, supporting the necessity of Aldh1l1 for FA rescue.

**Figure 3.**
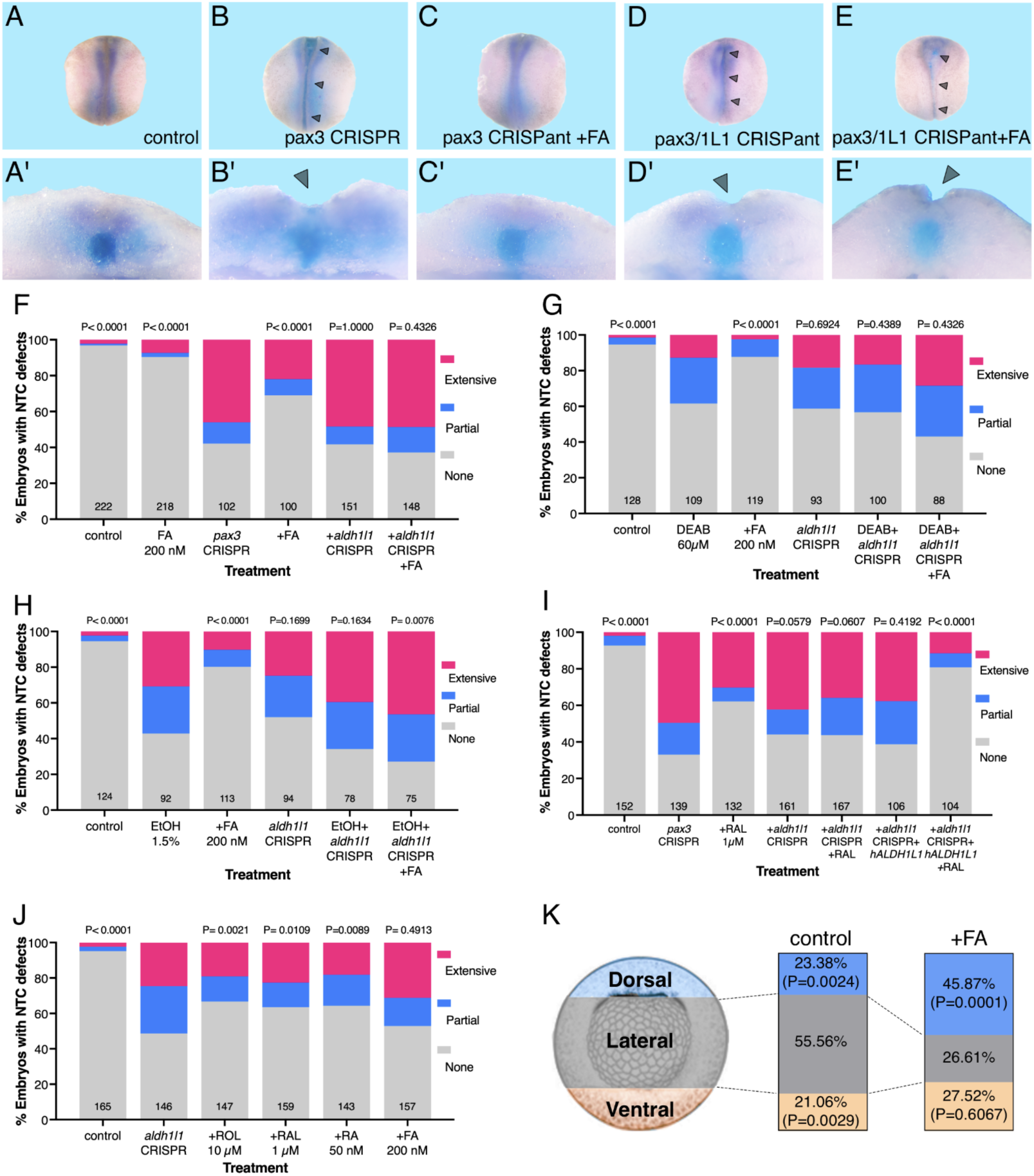
ALDH1L1 is required for FA rescue of NTC defects in *pax3* CRISPants and reduced RA embryos. In embryos treated with EtOH (1.5%), DEAB (60 µM) or *pax3* CRISPants, *aldh1l1* was targeted (CRISPR/Cas9) and supplemented with FA (200 nM) or RAL (1 µM), or injected with the human *hALDH1L1* RNA to rescue the NTC defects (arrowheads). (A-E’) CRISPants for *pax3* and *aldh1l1* were analyzed by *in situ* hybridization using probes for *sox3* to label the neural plate border (magenta) and *chrd* for the notochord (dark blue). (A-E) Dorsal views of embryos at NTC closure stages (NF19/20). (A’-E’) Detail of a cross-section of the same embryos bisected to demonstrate open neural tubes (B’, D’, E’). (F) NTC defect rescue analysis of *pax3* and *aldh1l1* double CRISPant embryos supplemented with FA. (G, H) The *aldh1l1* gene was targeted in embryos treated with DEAB (G) or EtOH (H) to induce NTC defects and study the rescuing effect of FA supplementation. (I) Rescue of NTC defects in *pax3* and *aldh1l1* double CRISPant embryos overexpressing hALDH1L1 and supplemented with RAL. (J) Rescue of NTC defects in *aldh1l1* CRISPant embryos was studied by supplementation with RA (50 nM), ROL (10 µM), RAL (1 µM), or FA (200 nM). The incidence was calculated as the fraction of embryos with NTC defects from the total sample. Sample sizes are shown, and statistical significance was calculated using Fisher’s exact test relative to the *pax3* CRISPant sample. (K) Changes in the *aldh1l1* transcript distribution induced by FA. Control and FA-treated gastrula stage (NF10.5) embryos were dissected into dorsal, lateral, and ventral marginal zone regions according to the scheme. qPCR for *aldh1l1* was performed, and the percentage of total transcripts in each region was calculated relative to whole embryos. Statistical significance was calculated relative to the lateral marginal zone abundance using one-way ANOVA.

The role of Aldh1l1 in the FA rescue of NTC defects induced by reduced RA signaling was also studied using our biochemically defined model. Embryos treated with DEAB (60 µM) or EtOH (1.5%) to inhibit RA production were made CRISPant for *aldh1l1* and supplemented with FA. DEAB does not inhibit the ALDH activity of Aldh1l1 (23). Knockdown of *aldh1l1* in DEAB-treated embryos resulted in a very slight, nonsignificant increase in NTC defects. FA supplementation was unable to reduce the incidence of NTC defects (Fig. 3G). Similar results were observed in EtOH (1.5%)-treated *aldh1l1* CRISPant embryos, which also exhibited a slight but nonsignificant increase in NTC defects compared to EtOH-only embryos (Fig. 3H). Also in this model, targeting *aldh1l1* prevented FA from rescuing the NTC defects (Fig. 3H), indicating again that Aldh1l1 activity is necessary for FA-driven rescue.

To demonstrate the necessity of Aldh1l1 for RA production and prevent NTC defects, we conducted rescue experiments overexpressing the human ALDH1L1 (hALDH1L1)(Fig. 3I). In *pax3* CRISPants, RAL supplementation enhances RA biosynthesis, effectively reducing NTC defects (Figs. 1F and 3I). Knockdown of *aldh1l1* hindered the RAL rescue of NTC defects, consistent with the loss of the ALDH activity required to oxidize RAL to RA (Fig. 3I). Injecting *hALDH1L1* mRNA into *pax3*+*aldh1l1* double CRISPants did not change the closure defects, while RAL supplementation in the presence of hALDH1L1 promoted efficient NTC defect rescue (Fig. 3I), reinforcing the observation that Aldh1l1 activity is necessary for FA to rescue NTC defects using RAL as a substrate.

To strengthen the link between Aldh1l1 and RA signaling, we investigated whether reduced Aldh1l1 activity could be rescued by providing ROL, RAL, RA, or FA. Supplementing *aldh1l1* CRISPant embryos with RA or its precursors, ROL or RAL, significantly reduced NTC defects (Fig. 3J). This suggests that other RALDH enzymes or residual Aldh1l1 activity might produce RA to rescue the defects. Importantly, FA supplementation did not rescue the defects (Fig. 3J), supporting the requirement for Aldh1l1 activity in FA rescue of neural tube closure malformations.

### Conservation of the FA–ALDH1L1–RA Pathway in Mammals

To biochemically demonstrate that ALDH1L1 can produce RA, we performed an enzymatic kinetic analysis of the human enzyme expressed in bacteria (17). The 100-kDa recombinant hALDH1L1 was detected in bacterial extracts using anti-His-tag antibodies (Supplementary Fig. 5), in agreement with its calculated size of 98.8 kDa. The affinity-purified hALDH1L1 oxidized all-*trans*-RAL (atRAL) to RA using NADP^+^ as a cofactor (19). The enzyme exhibited robust, concentration-dependent atRAL oxidizing activity (Fig. 4A). We determined the kinetic parameters of hALDH1L1 atRAL oxidation (Fig. 4B), A Km of 4.1 ± 0.47 µM and a Vmax of 2.94 ± 0.07 µmol/min per mg of enzyme were determined for ALDH1L1. These findings strongly support the hypothesis that FA promotes the upregulation of *ALDH1L1,* leading to localized RA production to prevent NTC defects.

**Figure 4.**
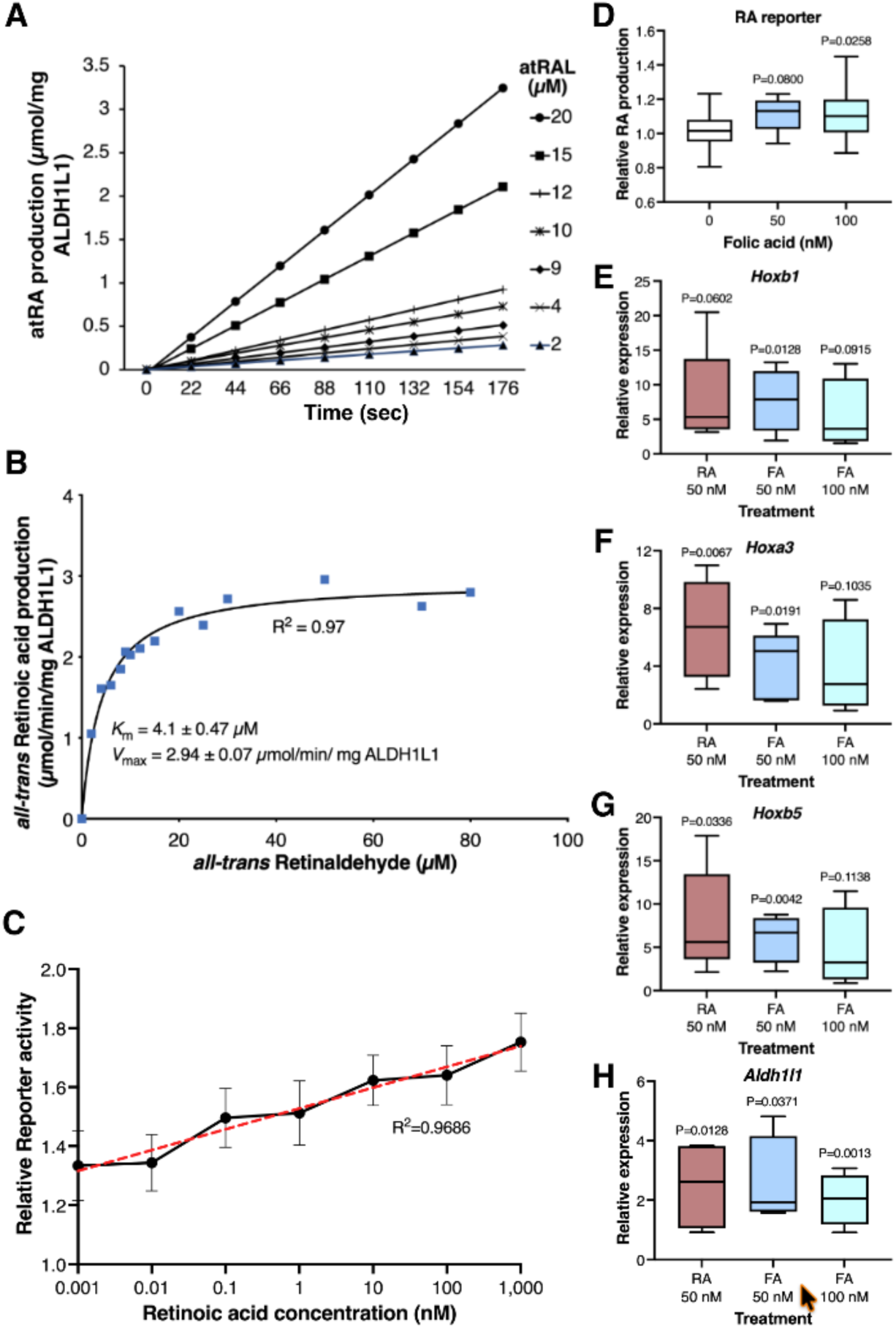
Human and mouse ALDH1L1 oxidize RAL to RA. The human ALDH1L1 protein was produced in bacteria as a His-tagged protein and then affinity-purified. (A) Production of all*-trans* RA *in vitro* at different substrate concentrations. (B) The human ALDH1L1 protein was subjected to enzymatic kinetic analysis for the production of all*-trans* RA by oxidation of atRAL in the presence of NADP^+^. Curve fitting was performed for the Michaelis-Menten constant calculation (*K*_m_). (C) Mouse F9 teratocarcinoma cells were stably transfected with the RA-reporter plasmid, pRAREhsplacZ. RA reporter cells were exposed to 1 pM to 1 µM RA. The relative reporter activity was calculated relative to the background activity in all experiments (N=7). The coefficient of determination (R2) for the line fit is shown. (D-H) RA reporter F9 cells were treated with 50 nM and 100 nM FA (N=5). The relative level of RA reporter activity (D), and the upregulation (qPCR) of RA-responsive genes, including *Hoxb1* (E), *Hoxb3* (F), and *Hoxb5* (G), and the *Aldh1l1* gene (H), was determined relative to controls. Box-and-whisker plots (5-95 percentiles) and statistical significance was calculated using one-way ANOVA compared to the control expression.

To demonstrate conservation of the molecular pathway uncovered in *Xenopus* embryos in mammals, we established an RA reporter cell system (27). F9 murine teratocarcinoma cells were stably transfected with the RA reporter plasmid, pRAREhsplacZ (28). RA reporter cells were exposed to all-*trans* RA concentrations from 1 µM to 1 pM to determine their RA responsiveness. They exhibited a dose-dependent response across the entire tested range (Fig. 4C). To investigate whether FA promotes RA production in mouse cells, reporter cells were treated with FA (50 nM and 100 nM), and reporter activity was measured. The RA reporter cells showed a proportional response to FA addition (Fig. 4D). The RA-responsive genes *Hoxb1*, *Hoxa3*, and *Hoxb5* were also upregulated by FA in RA reporter cells (Fig. 4E-G). Significantly, *Aldh1l1* was also upregulated in mouse reporter cells treated with FA (Fig. 4H). These results show that in mouse cells, FA also increases RA signaling and upregulates *Aldh1l1* expression.

### Folic Acid Alters the Spatial Expression of ALDH1L1 During Gastrulation

To gain deeper insight into the effects of FA exposure, we investigated whether, in addition to the up-regulation, the spatial expression pattern of *aldh1l1* is also altered as part of the self-regulatory response of FA metabolism (Fig. 2M and Supplementary Fig. 3E). Analysis of *aldh1l1* expression from blastula (NF8) to NTC stages (NF19) showed that transcripts begin to accumulate close to the onset of gastrulation (NF10) and increase as gastrulation progresses (Supplementary Fig. 6). This expression pattern aligns with the appropriate developmental window when FA efficiently promotes closure in both NTC defect models studied (Figs. 2A-C).

The *aldh1l1* spatial expression pattern was analyzed by dissecting embryos into different regions at early/mid gastrula (NF10.5), early neurula (NF15), and late neurula (NF19) stages. During early/mid gastrula, relative transcript abundance was determined comparing dorsal, lateral, and ventral marginal zone regions (Figs. 3K and Supplementary Fig. 7A). At this stage, most *aldh1l1* transcripts localize to the lateral region (LMZ), with limited expression in the dorsal (DMZ) or ventral (VMZ) regions (Figs. 3K and Supplementary Fig. 7A). To confirm the dissections, we examined *chrd*, *myod1*, and *szl* as markers for the DMZ, LMZ, and VMZ, respectively (Supplementary Fig. 7B-D). The *aldh1l1* transcripts co-localized with *myod1* expression in the more lateral regions.

During early neurula (NF15), embryos were dissected into neural tube (NT) and non-neural (nonNT) regions, with *aldh1l1* transcripts being more abundant in nonNT (Fig. S7I). Dissections were confirmed with *sox3* and *pax3* for NT, and *pax8* for nonNT (Supplementary Fig. 7J-L). During NTC stages (NF19), embryos were dissected into NT and nonNT. In some, the NT was further divided into cranial (crNT) and trunk (trNT), with *aldh1l1* more abundant in nonNT (Supplementary Fig. 7M). Dissection accuracy was validated with *pax3* for NT and crNT, *otx2* for cranial NT, and *hoxc6* for trunk NT (Supplementary Fig. 7N-P). These results show that *aldh1l1* remains localized to lateral, non-neural regions.

As FA most efficiently rescues NTC defects during early gastrulation (Fig. 2A-C), the effect on *aldh1l1* expression was studied in dissected regions from FA-supplemented gastrula embryos. FA treatment increases *aldh1l1* expression by approximately 1.5-fold (Figs. 2M, Supplementary Figs. 3D, and 7E). However, FA treatment also altered the spatial distribution of *aldh1l1* transcripts. The DMZ showed a significant increase in relative *aldh1l1* transcript abundance from 23.38% to 45.87% (Figs. 3K and Supplementary Fig. 7F). The LMZ decreased from 55.56% to 26.61%, and the VMZ slightly increased (Figs. 3K and Supplementary Fig. 7G, H). The enhanced *aldh1l1* expression in the DMZ places the FA-increased Aldh1l1 activity near the forming neural plate (8).

### ALDH1L1 Activity Restricts Neural Plate Expansion During FA Rescue

Reduced RA levels during early gastrulation increase neural precursor cell proliferation, leading to a teratogenic expansion of the neural plate and NTC defects (8). To investigate the role of *aldh1l1* in this early phenotype, we conducted experiments in which one side of the embryo was CRISPant for *aldh1l1*. FITC-dextran was co-injected with the Cas9/sgRNA riboprotein to label the CRISPant side (8). Embryos were treated with DEAB to induce NTC defects, and FA was added for defect rescue (Fig. 5A). Proliferating cells in the neural plate were labeled with antibodies against phosphorylated histone H3 (pHH3) (Fig. 5B-E)(8). Control embryos were injected with FITC-dextran on one side (Fig. 5B) or uninjected. The number of proliferating cells in the neural plate was determined in the right and left halves of the embryo. One-sided targeting of *aldh1l1* resulted in a slight but statistically significant increase in proliferation on the CRISPant side compared to the control side (Fig. 5C). Cell proliferation rose to an average of 59.5 pHH3-positive cells on the CRISPant side from an average of 51.8 pHH3-positive cells on the control side (Fig. 5F; Δ=7.7, n=38). The DEAB treatment increased proliferation in the neural plate (63.5 pHH3-positive cells), exhibiting a weak additive effect with *aldh1l1* knockdown (69.9 pHH3-positive cells), as the treatment is systemic (Figs. 5D, F; Δ=6.4, n=46). FA supplementation restored normal proliferation levels in the control side of DEAB-treated embryos (52.6 pHH3-positive cells), while the *aldh1l1* CRISPant side still exhibited increased proliferation (Figs. 5E, F; Δ=28.9, n=42). Control FITC-dextran injected sides showed no difference compared to the control uninjected side (Figs. 5B, F; Δ=-0.14, n=51). Then, inhibition of RA biosynthesis (DEAB) induced the previously observed increase in cell proliferation, which can be rescued by adding FA, but it requires a functional *aldh1l1* gene.

**Figure 5.**
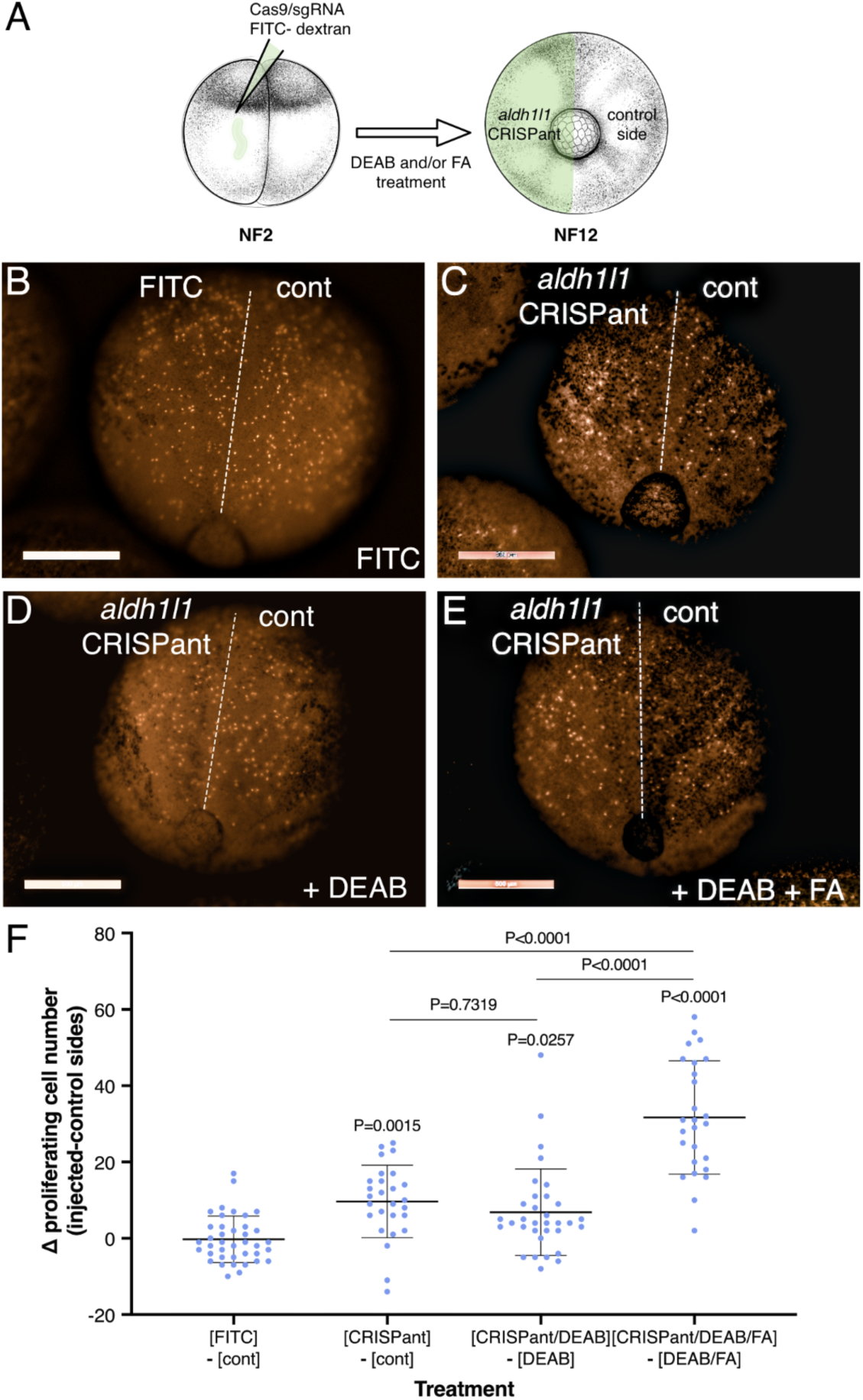
The ALDH1L1 activity is required for FA to restore normal neural plate proliferation. (A) Description of the experimental design to analyze the ALDH1L1 activity in rescuing cell proliferation in the neural plate. (B-F) Embryos were injected with Cas9/sgRNA riboprotein and FITC-dextran in one blastomere at the two-cell stage and stained for phosphorylated histone H3 to monitor cell proliferation. (B) Embryo injected with FITC-dextran on one side. (C) One-sided *aldh1l1* CRISPant embryo. (D) One-sided *aldh1l1* CRISPant treated with DEAB to induce NTC defects. (E) Treatment of a one-sided *aldh1l1* CRISPant with DEAB and FA. (F) The number of phospho-histone H3 positive cells was determined on each embryo side, and the difference between the CRISPant and control sides in each embryo was calculated. Each dot represents one embryo, and statistical significance was calculated using one-way ANOVA.

### Optimizing Folic Acid and Retinoid Supplhementation for NTD Prevention

The results described reveal a novel link between FA supplementation and localized RA biosynthesis, which explains how FA prevents NTC defects. We directly tested whether retinoid supplementation improves FA rescue in embryos with reduced RA (DEAB) (Figs. 1L, M). Low FA treatment (25 nM instead of 200 nM) weakly rescued NTC defects (Fig. 6B). Adding RA (20 nM or 5 nM) significantly improved the rescue efficiency (Fig. 6B). Higher RA levels (50 nM) on their own can rescue NTC defects (8), as it bypasses the need for the retinaldehyde dehydrogenase activity, endogenous or FA induced (Fig. 6A). To link FA upregulation of ALDH1L1 and physiological RA production, DEAB-treated embryos were supplemented with low ROL (vitamin A) concentrations (5 µM and 1 µM) instead of 10 µM that rescues NTC defects (8). Normally, ROL is oxidized to RAL, the ALDH1L1 substrate (Figs. 4A, B, and 6A). Low ROL or FA (50 nM) individual treatments resulted in weak rescues, insufficient to prevent NTC defects (Fig. 6C). However, combined FA (50 nM) and ROL (5 µM) treatment led to very efficient rescue, improving upon FA alone (Fig. 6C). These results show that vitamin A within the physiological range enhances FA rescue.

**Figure 6.**
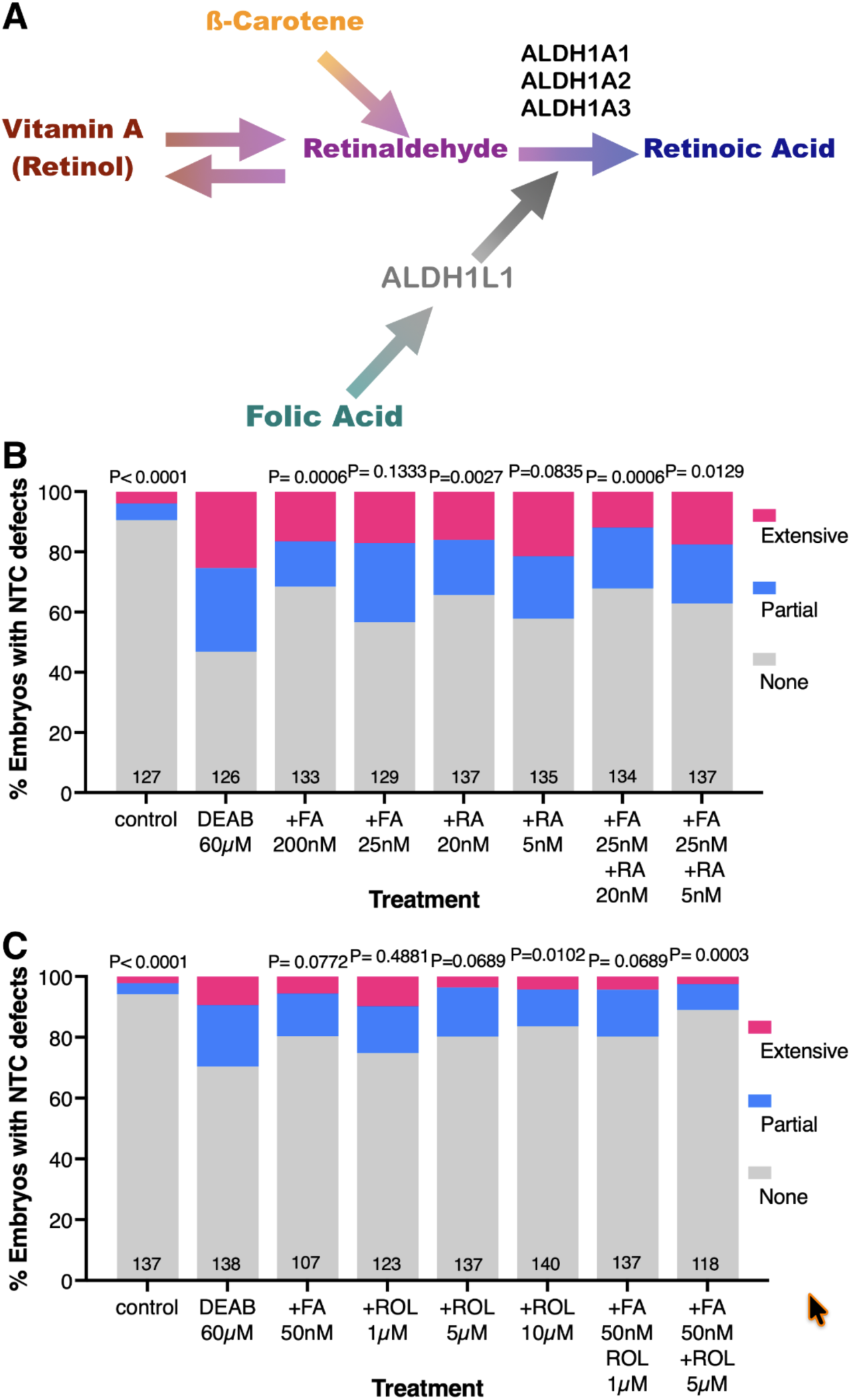
Retinol improves the rescue efficiency of NTC defects by FA. (A) Schematic representation of the RA biosynthetic pathway and the proposed role of FA and ALDH1L1 in RA production. The main RA precursors are shown, as well as the main retinaldehyde dehydrogenases (ALDH1A1/2/3). (B, C) Embryos were treated with DEAB to induce NTC defects, and rescue experiments were performed with low FA concentrations (50 nM or 25 nM) and low amounts of (B) RA (20 nM or 5 nM), or (C) ROL (vitamin A; 1 µM-10 µM). The incidence was calculated as the fraction of embryos with NTC defects from the total sample. Sample sizes are shown, and statistical significance was calculated using Fisher’s exact test relative to the *pax3* CRISPant sample.

## DISCUSSION

### FA supplementation increases RA levels

We previously induced NTC defects by reducing RA levels using pharmacological inhibitors of retinaldehyde dehydrogenases, and later rescued them with RA precursor supplementation (8). Now, we show that FA supplementation also rescues RA-induced NTC defects via RA production, as evidenced by upregulation of RA-responsive genes (*Hox* and RA metabolism) (17) within a similar developmental rescue window (8). Reduced RA signaling is crucial in FASD etiology (13, 16, 17, 29, 30), and FA can reverse developmental malformations in FASD or decreased RA (30–37). The FA ability to rescue NTDs and alcohol-induced malformations reveals a new biochemical connection between one-carbon (folate) metabolism and RA signaling.

The *Splotch* (*Pax3*) mutant mouse is a well-studied model for FA rescue of NTDs (38, 39). *PAX3,* mutated in Waardenburg syndrome, sometimes causes NTDs (40). CRISPR analysis of *pax3* NTDs showed that FA treatments work within the same sensitivity window as those rescuing NTDs caused by reduced RA (8). Retinoid supplements also rescued NTDs in *Xenopus pax3* CRISPants, similar to the mouse *Splotch* model (24). These findings suggest that FA rescues NTDs in reduced RA or *pax3* CRISPants involving RA production.

### The ALDH1L1 activity is required to rescue NTC defects with FA

Multiple models explain how FA prevents NTDs, involving DNA repair, methylation, neurodevelopment, epigenetics, nucleotide biosynthesis, and cell proliferation (39, 41, 42). However, these fail to clarify the biochemical connection, the timing, gene-environment interactions, or the selective rescue observed in animal models and humans. No detailed molecular pathway has been proposed for how FA prevents NTDs during neural tube formation. The folate metabolic network is regulated at transcriptional, post-translational, allosteric, and cofactor levels to balance one-carbon donors (43), affecting gene expression and epigenetics (44). We found that FA upregulates the one-carbon metabolism gene *aldh1l1,* which encodes ALDH1L1 and has homology to retinaldehyde dehydrogenases (19, 20). FA also increased the expression of RA target genes. Using a murine RA-reporter cell line, we demonstrate conservation of FA-induced *Aldh1l1* upregulation and RA production in mammals.

Rescue of NTC defects with FA in *Xsplotch* or reduced RA failed if embryos were CRISPant for *aldh1l1*, showing that ALDH1L1 is essential for FA to rescue NTC defects. Human *ALDH1L1* overexpression restored FA rescue in *Xsplotch* CRISPant embryos, and this effect was further improved by RAL, indicating that ALDH1L1 produces RA to prevent NTDs. Human ALDH1L1 oxidizes all-*trans* RAL with a Km of 4.1 µM, which falls within the range of previously reported constants (0.2 µM - 16 µM) for retinaldehyde dehydrogenases (45), supporting the FA-RA link. In humans, *ALDH1L1* polymorphisms were associated with NTD induction, further supporting the role of ALDH1L1 in rescuing NTC defects induced by FA.

We previously showed that early effects of reduced RA signaling include upregulation of the neural precursor regulators *foxd4l1.1* and *gmnn* (8). Neural induction activates these factors that delay differentiation and promote neural precursor proliferation (46). Reduced RA increases neural precursor proliferation and enlarges the neural plate (8). FA supplementation rescues overproliferation, which is prevented in *aldh1l1* CRISPants, showing that ALDH1L1 is needed for FA to rescue neuroectodermal proliferation. Spatial expression analysis revealed that FA upregulates and dorsalizes *aldh1l1* expression, indicating that FA alters *aldh1l1* transcript levels and localization, promoting RA production by ALDH1L1 near the neuroectoderm and reducing NTDs.

### Vitamin A improves the efficiency of FA in preventing NTDs

The upregulation of *aldh1l1* after FA exposure and increased RA signaling offer a biochemical explanation for how FA helps prevent NTDs. ALDH1L1 needs RAL, derived from vitamin A oxidation, to produce RA. We tested whether combined vitamin A and FA (vitamin B9) improved NTD rescue. Reducing FA supplementation (25-50 nM) to serum levels (7-45 nM) resulted in weak rescue of RA-induced NTDs. Adding small amounts of RA improved rescue, as it bypasses RALDH. Combining low ROL with low FA significantly rescued NTDs, whereas low ROL alone did not. These results show that ROL enhances the efficiency of FA prevention of NTDs, with FA upregulating ALDH1L1 to produce RA from RAL.

Vitamin A can sometimes reach teratogenic levels from diet, supplements, and acne medications (47). Excess vitamin A causes birth defects in humans, such as *spina bifida*, while animal studies show that moderate RA levels can prevent NTDs. Maintaining RA within a narrow range is crucial for neural tube closure (4, 48). Human genetic studies have linked NTD risk to polymorphisms in RA metabolism genes, some of which may reduce RA signaling (49). Typical FA supplementation doses (0.4-0.6 mg/day) reduce NTD risk by about 50% (50), depending on genetics (41). Higher doses (up to 5 mg/day) are prescribed for at-risk women, including those with a history of alcoholism. Recent research recommends increasing FA to nearly 100 nM in serum to improve NTD prevention (50), despite concerns about its teratogenic effects (11, 12). Our findings reveal a metabolic mechanism underlying the protective effect of FA against abnormal neural tube closure, and provide initial evidence that RA can enhance the efficiency of FA in preventing NTDs. Moreover, our results may explain cases of FA-unresponsive pregnancies, as arising from impaired RA production by ALDH1L1.

## MATERIALS and METHODS

### Embryos and treatments

*Xenopus laevis* frogs were purchased from Xenopus 1 or Nasco (Dexter, MI or Fort Atkinson, WI, United States). Experiments were performed after approval and under the supervision of the Institutional Animal Care and Use Committee (IACUC) of the Hebrew University (Ethics approval no. MD-21-16767-3). Embryos were obtained by *in vitro* fertilization, incubated in 0.1% Modified Barth’s Solution and HEPES (MBSH), and staged according to Nieuwkoop and Faber (NF) (NF; 51). Embryos were treated with the RA signaling inhibitors: ethanol (0.5% and 1.5% vol/vol; Bio-Lab 000525250200) and 4-Diethylaminobenzaldehyde (DEAB, 60 µM, stock dissolved in DMSO, D86256 Sigma-Aldrich) or with *all-trans* Retinol (ROL, 10 µM, R7632 Sigma-Aldrich), *all-trans* Retinaldehyde (RAL, 1 µM, R2500 Sigma-Aldrich), and *all-trans* Retinoic Acid (RA, 50 nM, R2625 Sigma-Aldrich). Folic Acid (F8758, Sigma-Aldrich) was prepared fresh by dissolving in 1M NaOH at 37°C, diluted to 100 mM stock, and further diluted in 0.1% MBSH to 500 µM at 37°C. Methotrexate hydrate (MTX, A6770, Sigma-Aldrich) was dissolved in 1M NaOH to 11 mM (5 mg/mL) and further diluted in 0.1% MBSH to 0.55 µM (0.25 µg/mL). Treatments were performed in 0.1% MBSH from the midblastula transition (MBT, NF8.5) until the desired analysis stage. For the temporal window determination experiments, embryo treatments were initiated at different stages (NF8, NF10.25, and NF11). Embryos of both sexes were analyzed throughout this project, as at the stages studied, they are indistinguishable morphologically.

### Embryo manipulation and dissection

Embryos were dissected into three regions at NF10.5: dorsal, lateral, and ventral marginal zone; Two regions at NF15: neural tube (NT) and non-neural tube (non-NT); and four embryonic regions at NF19: neural tube (NT), cranial neural tube (crNT), trunk neural tube (trNT), and non-neural tube (non-NT). Embryos were dissected in 1X MBSH buffer. Embryonic regions (15-20 embryos for each region) were then processed for RNA extraction and real-time RT-PCR.

### Gene targeting with CRISPR\Cas9

For gene-specific targeting with CRISPR/Cas9, single guide RNAs (sgRNA) were designed using genomic DNA sequences selected from Xenbase.org (52) for the L and S homoeologs when present. We used CRISPRdirect (53) and CRISPRscan (54) for target site search. Computational estimation of the sgRNA efficiency was determined using the inDelphi software (55, 56). For the generation of F0 CRISPant embryos, we injected one-cell stage embryos with Cas9 ribonucleoprotein (RNP) complexes employing the two-RNA component (crRNA:tracrRNA) approach (57). The sgRNA targeting *pax3.*L/S is 5’GGCAGGGUGAACCAGCUGGG; to target *aldh1l1,* we used 5’CCUUACUGCUGGCAACACAG. Briefly, chemically synthesized and modified for stability (Alt-R) RNAs (crRNA and tracrRNA; IDT, United States) were annealed to generate the double guide complexes (crRNA:tracrRNA) and were incubated (10 min at 37°C) with *S. pyogenes* Cas9 protein (IDT, United States) to generate RNP complexes. Eight nanoliters of the RNP complex solution were injected into the cytoplasm of one-cell stage *Xenopus* embryos. For the proliferation analysis, guide RNA was injected into one cell at the 2-cell stage (NF2) with FITC-dextran (70 KDa, 50mg/ml stock diluted 1:32-64, FD70 Sigma-Aldrich) to detect the injected side; these embryos were fixed at the desired stage for analysis. To determine the efficiency of Indel induction, genomic DNA was extracted from 5 individual embryos at neurula (NF19-20), employing the GenElute Mammalian Genomic DNA Miniprep Kit (SIGMA). The genomic region containing the CRISPR/Cas9 targeted region was PCR amplified using a nested PCR approach (Table 1), and the size-selected and cleaned product was sequenced. Genome editing efficiency was analyzed by decomposition analysis using the Synthego ICE algorithm (58) or TIDE (22).

**Table 1.**
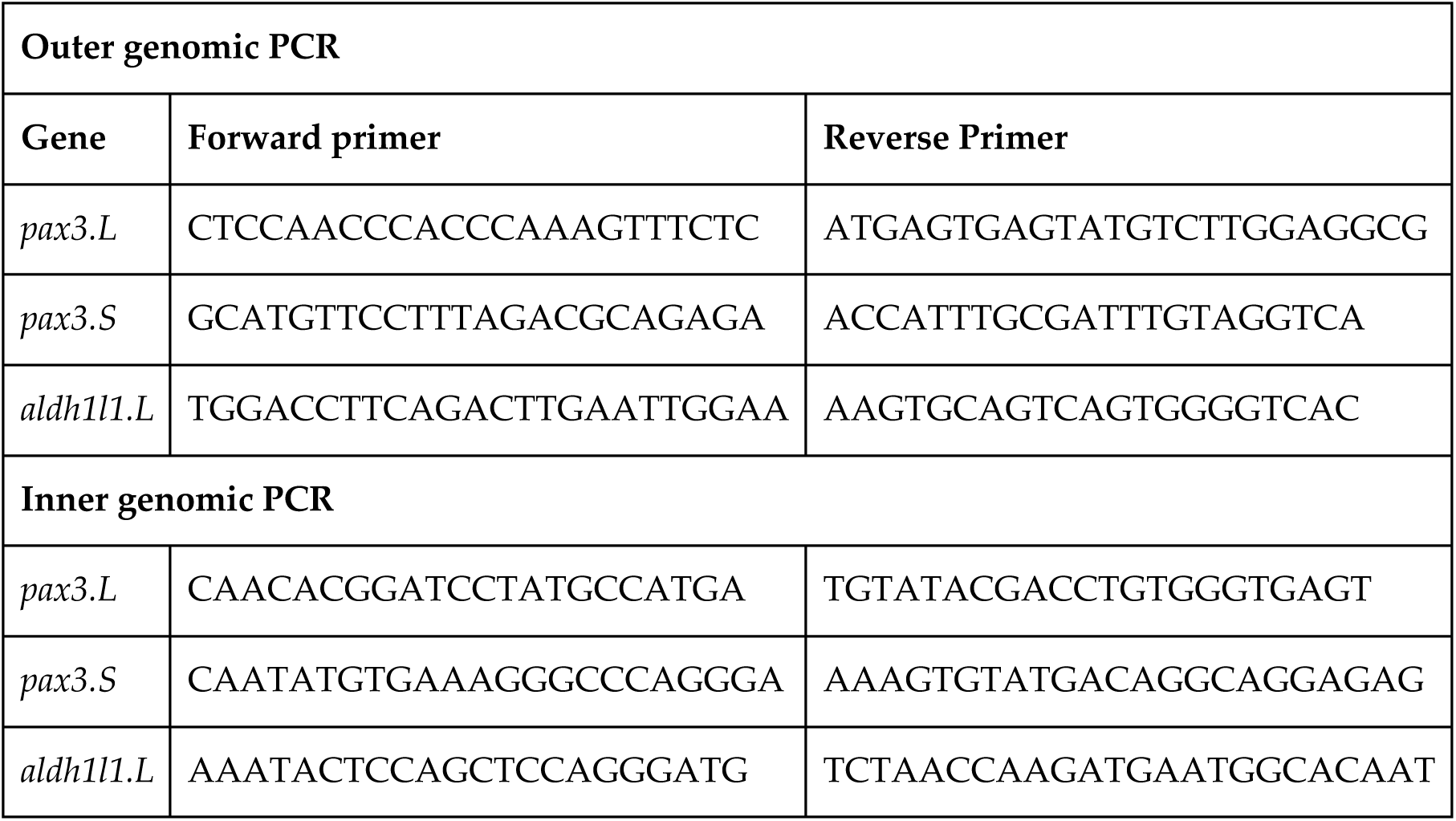
Primers for estimation of the CRISPR/Cas9 gene targeting efficiency.

### RNA injection

Embryos were injected with *in vitro* transcribed capped mRNA. Capped mRNA of the *cyp26a1* gene was prepared from the pCS2 NotI linearized plasmid using SP6 RNA polymerase (59). Capped mRNA of the human *aldh1l1* gene (Source Bioscience) was prepared from a pCS2 linearized plasmid, cut with NotI, using SP6 RNA polymerase. Cap analog (m7G (5′)ppp (5′)G; New England Biolabs, USA) was added to the reaction mixture using a cap:GTP ratio of 5:1. To analyze the injection site, mRNA solutions were mixed with FITC-dextran (70 KDa, 50mg/ml stock diluted 1:32, FD70 Sigma-Aldrich).

### NTC defect analysis

The frequency of NTC defects was evaluated following a morphological examination of neural tube closure integrity at late neurula (NF19-20). All embryos and samples were kept at the same temperature and conditions throughout the experiments. The embryos were divided into four groups by the location and severity of the NTC defects. The embryos were counted and fixed for validation by *in situ* hybridization and bisection. The NTC defects analysis was performed independently by two different researchers.

Whole-mount and double *in situ* hybridization were performed as previously described (8, 60). Embryos were fixed at NF19-20 in MEMFA and processed for whole-mount *in situ* hybridization. Probes were prepared by *in vitro* transcription using Digoxigenin- or Fluorescein-labeled nucleotide mixes (11277073910 and 11685619910, Roche). Probes were transcribed as previously described: *sox3* (*61*), *chrd.1* (*chordin*) (62), *pax3* (63), and *shh* (64).

### Analysis of cell proliferation

Immunofluorescence staining was performed on whole-mount embryos. At the onset of gastrulation, embryos were separated into groups based on the injection side (right or left) based on the FITC-dextran, then incubated until late gastrula and fixed for 1–2 hours at room temperature in MEMFA (100 mM MOPS (pH 7.4), 2 mM EGTA, 1 mM MgSO_4_, 3.7% (v/v) formaldehyde). Embryos were then permeabilized with PBS 1X with 0.1% Triton-X100 and 0.2% BSA solution, incubated in blocking solution (Cas-Block; LIFE technologies; 008120), and stained according to standard procedures. Primary antibody: mouse anti-phospho-Histone H3 (Ser10; 1:200; cat# 9706; Cell Signaling Technology). Secondary antibody: Rhodamine 590-conjugated donkey anti-rabbit IgG (1:500, Jackson ImmunoResearch Laboratories, 711-295-152). The fluorescence analysis was performed using a Leica M165 FC fluorescence stereo microscope (Leica Microsystems). Image capture was performed using the Leica DFC3000 G monochrome camera, controlled by the Leica application suite (LAS) V4.12.

### Gene expression analysis by quantitative real-time RT-PCR

Total RNA from embryos was extracted with the Aurum Total RNA Mini Kit (7326820 Bio-Rad), and cDNA was synthesized using iScript cDNA Synthesis Kit (95047-100 QuantaBio). The real-time PCR reactions were performed using the CFX384 Real-Time System (Bio-Rad) and iTaq Universal SYBR Green Supermix (1725124 Bio-Rad). Each experiment was repeated at least three times independently, and the samples were run in triplicate each time. *slc35b1*.L was the housekeeping reference gene (65). The primers used for qPCR analysis are listed in Table 2.

**Table 2.**
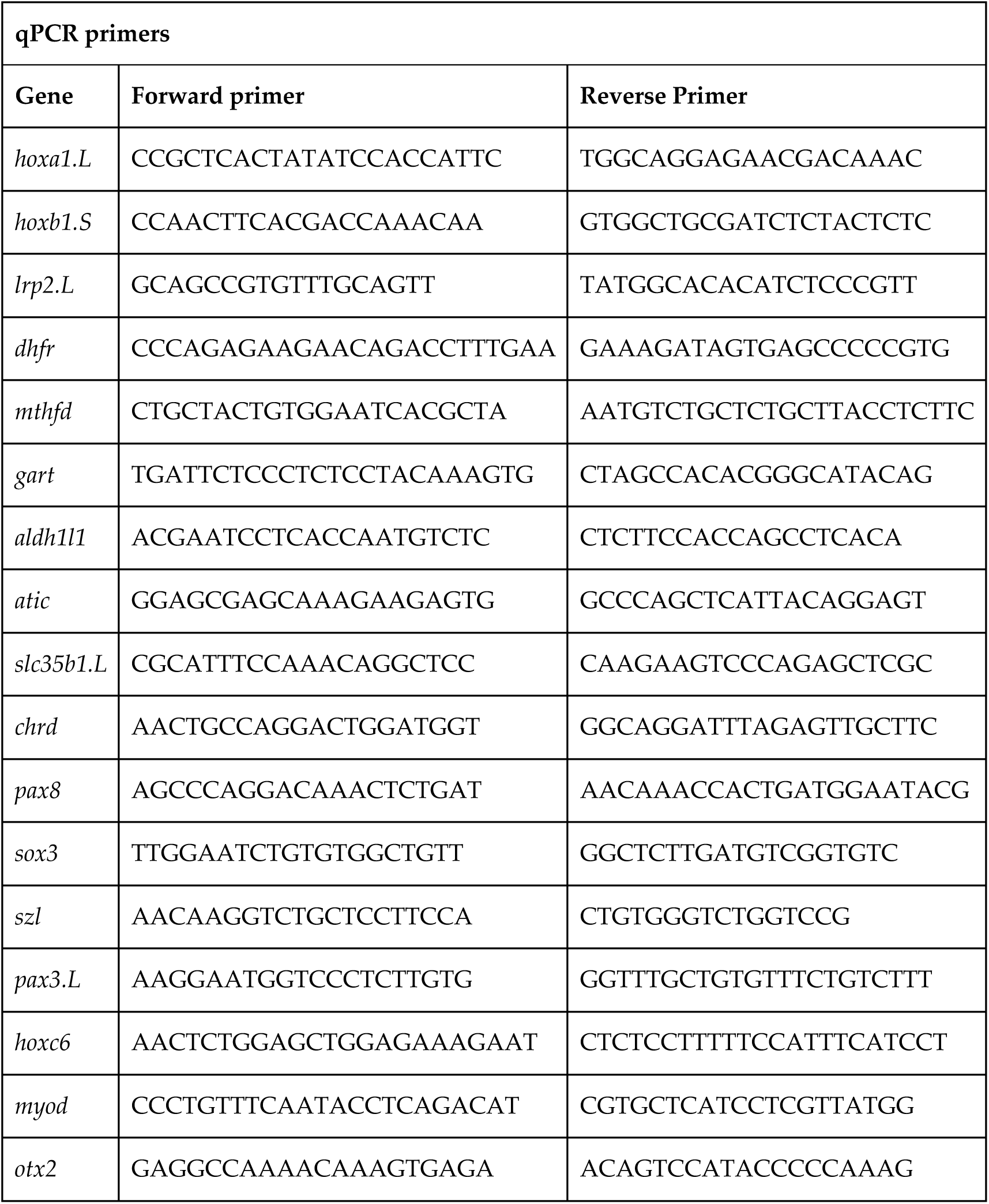
Primers for PCR expression analysis.

### Statistics

Statistical significance calculations were done using the Prism software package (GraphPad Software Inc., San Diego, CA). Tests used were ANOVA and t-test, depending on the type of data analyzed. Differences between means were considered significant at a significance level of p < 0.05. Fisher’s Exact Test was performed using the GraphPad site or VassarStats (vassarstats.net).

### Kinetic analysis of the RA-producing activity of ALDH1L1

For recombinant protein production, the human *ALDH1L1* cDNA was BamHI and ClaI cloned in the pET28a (Novagen) vector for expression and His-tag purification. The ALDH1L1 protein was expressed in *E. coli* strain BL- 21 CodonPlus after isopropyl β- D -1-thiogalactopyranoside (IPTG) induction for 2 h at 37°C. Bacterial pellets were sonicated in 20 mM Tris-HCl (pH 7.4), 1 mM EDTA, 100 mM NaCl, imidazole, and complete protease inhibitor mixture (Roche Life Science). His-tagged ALDH1L1 was affinity-purified on Ni-NTA agarose resin (Qiagen). Partial ALDH1L1 protein fragments in the eluate were removed using an Amicon Ultra centrifugal filter 30 kDa (Millipore). The purity of the recombinant proteins was determined by SDS-polyacrylamide gel electrophoresis (12%), stained with Coomassie Brilliant blue, and western blotting to detect the his-tagged protein. We used the mouse anti-His antibody (GE Healthcare) at a 1:10,000 dilution. Immunoreactive proteins were detected using a chemiluminescence kit (Bet HaEmek Industries, Israel) according to the manufacturer’s protocol.

The ALDH1L1 enzymatic reaction was performed using the conditions previously described (66). Briefly, reactions were performed at 30°C in 50 mM Hepes/K^+^ pH 7.6 buffer containing 2 mM MgCl2, 2 mM DTT, 150 mM KCl, and 1 mM EDTA. Protein content in the reaction mixtures was 0.5µg/ml. Reactions were initiated by adding different concentrations of the *all-trans* retinaldehyde (1-80 µM) substrate and 1 mM NADP^+^ (10128031001 Sigma-Aldrich). Standard reactions were included in each experiment. Oxidation of retinaldehyde by ALDH1L1 was monitored by spectrophotometer determination (OD_340_) of the NADP^+^ reduction. The contribution of RAL absorbance was subtracted from the overall increase in absorbance (*A*) of the produced NADPH and RA. A global extinction coefficient of 15,900 M^−1^cm^−1^ was experimentally determined by measuring the absorbance change of increasing NADPH or *all-trans* RA concentrations under our reaction conditions (67). *V*_initial_ measurements were carried out at 30°C on a TECAN Infinite F200Pro spectrophotometer, sampling the absorbance every 10 sec for 30 min. The *V*_initial_ for each substrate concentration was calculated from the slope of the reaction kinetics in the linear range of the reaction. The synthesis rates (*V*_initial_) were then plotted as a function of substrate concentration, and *K*_m_ and *V*_max_ were obtained by nonlinear regression of the data to the Michaelis–Menten equations using Prism software (GraphPad Software, Inc., San Diego, CA).

### RA detection using RA reporter cells

To establish an RA reporter cell line (27), F9 cells were transfected with the RA reporter plasmid pRAREhsplacZ (28). The cells were co-transfected with the pHKO_09 plasmid based on the pLKO.1 puro plasmid to provide puromycin resistance (68). The Cells were transfected with Mirus (TransiT-LT1, Mirus-Bio, Merck) according to the manufacturer’s recommendations. Single clones were isolated and tested for RA responsiveness by titrating *all-trans* RA from 10 µM to 1 pM according to the procedure described by Lee et al. (27). To detect the RA produced *in vitro*, 10 µl of the enzymatic reaction were added to the well and incubated for 18 hours for analysis following Xgal staining and spectrophotometric quantification (27). An RA calibration curve was always run in parallel to the experimental samples. All samples were tested in triplicate for each biological replicate.

## DECLARATIONS

### Availability of data and materials

The original contributions presented in the study are included in the article and Supplementary Material; further inquiries can be directed to the corresponding author.

### Authors’ contributions

TE, TAL, and YS designed and performed the experiments, analyzed the data, and helped write the manuscript. JMI, DC, and GP performed some experiments and assisted with data analysis. AF and JAB supervised the project, designed experiments, analyzed the data, wrote the manuscript, and procured funding.

### Ethics approval and consent to participate

Animal experiments were performed after approval and under the supervision of the Institutional Animal Care and Use Committee (IACUC) of the Hebrew University (Ethics approval no. MD-21-16767-3).

## Acknowledgments

We thank Horst Grunz for the sox3 probe, Yaniv Elkouby for using his fluorescence microscope setup, and Sally Moody and Martin Blum for lengthy discussions, advice, and extensive comments on the manuscript.

