## Supplemental Figures for "Folic acid prevention of neural tube defects requires retinoic acid produced by ALDH1L1"

### SUPPLEMENTARY FIGURES

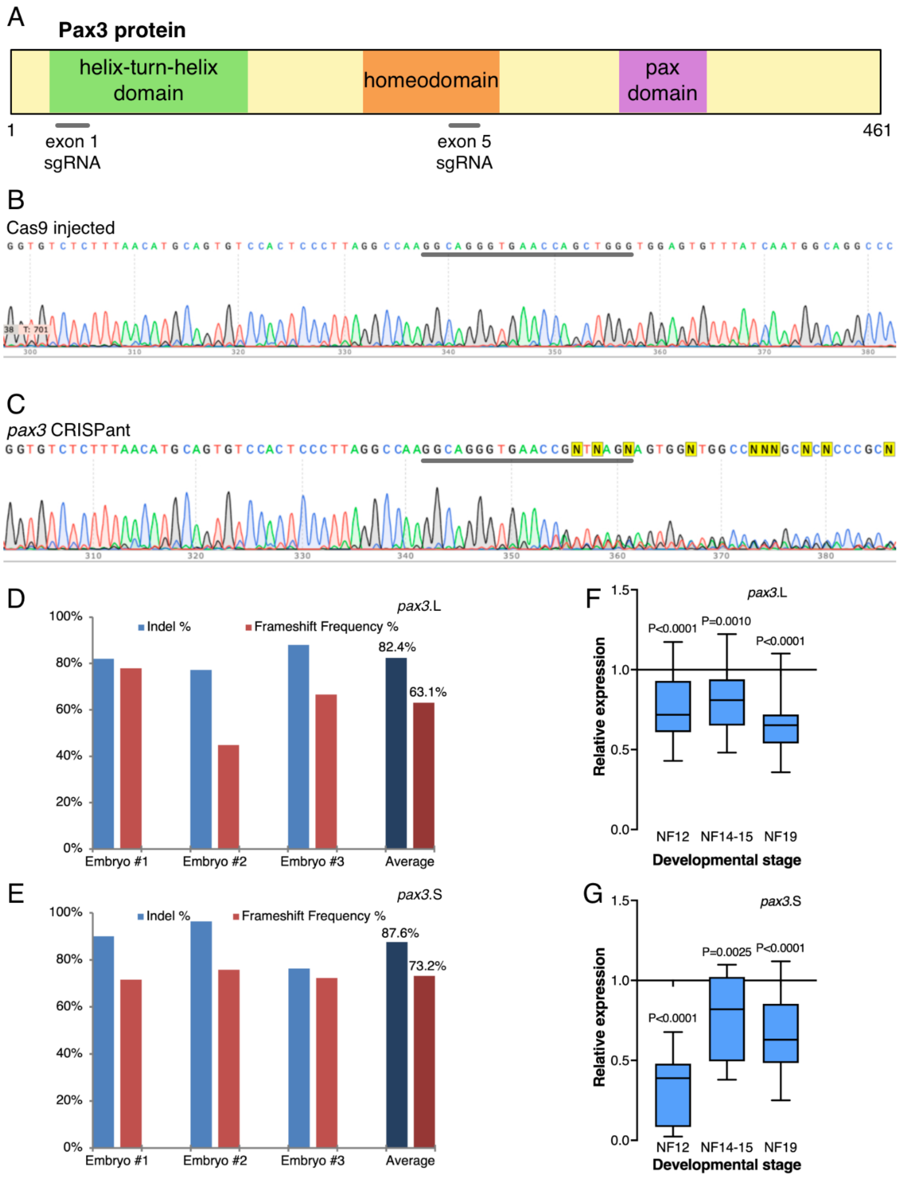


**Supplementary Figure 1. *Xsplotch, pax3* CRISPant embryos in *X. laevis*.** (A) Schematic representation of the Pax3 protein and the sgRNAs tested. (B, C) Sequencing traces of genomic DNA extracted from control Cas9-injected (B) and *pax3* CRISPant embryos at NF19-20 (C). The position of the sgRNA is shown (underlined). (D, E) TIDE analysis for indels and frameshift mutations in the *pax3* homoeologs (L and S) in individual CRISPant embryos. (F, G) Expression analysis (qPCR) of *pax3*.L and *pax3*.S in CRISPant embryos during late gastrula (NF12), early neurula (NF14-15), and NTC stages (NF19). Box and whiskers (5-95 percentile) plot, significance was calculated (ANOVA) relative to control embryos at the same stage.

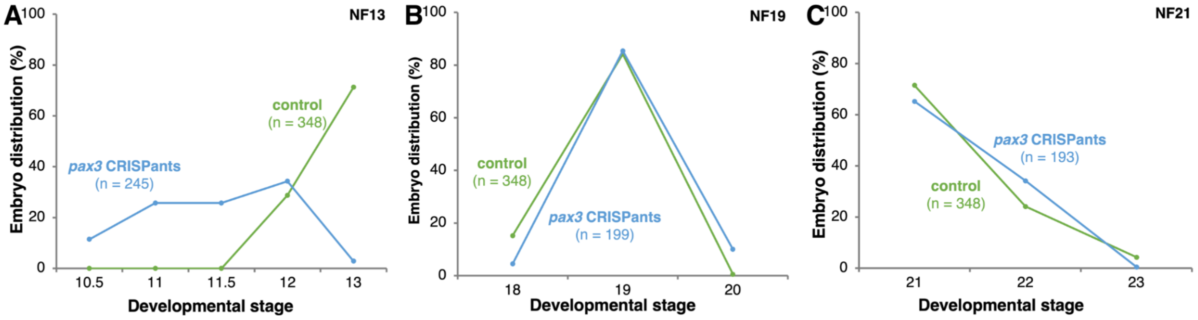


**Supplementary Figure 2. Transient developmental delay in *pax3* CRISPant embryos.** The same *Xsplotch* embryos were staged several times as they progressed through embryogenesis. All embryos were staged when the control siblings from the same batch reached NF13 (A), NF19 (B), or NF21 (C). The stage distribution of the embryos is shown as a percentage of the sample size.

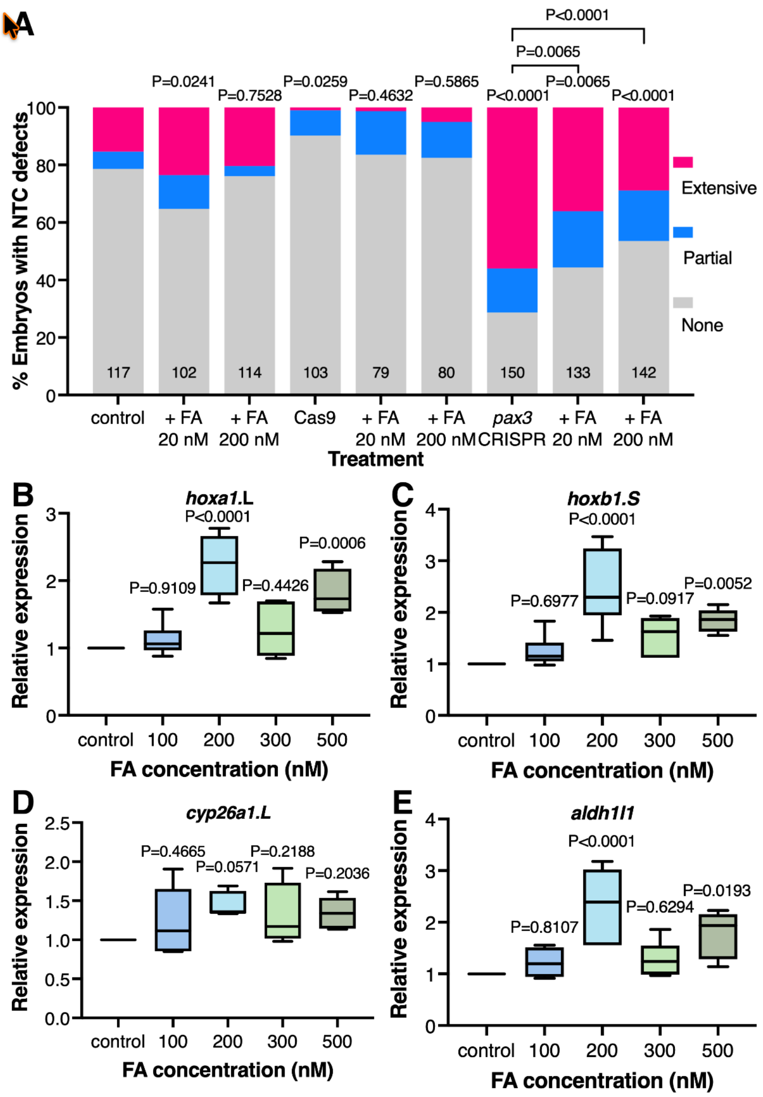


**Supplementary Figure 3. FA concentrations that up-regulate RA target genes.** (A) *Pax3* CRISPant embryos were treated with 20 nM or 200 nM FA to rescue the formation of NTC defects. Controls included uninjected embryos and embryos injected with Cas9 only. Embryos were scored at NF19 for NTC defects. The NTC defect incidence was normalized to the control sample. Sample sizes are shown, and statistical significance was determined using Fisher's exact test. (B-E) Embryos were treated with increasing FA concentrations (100 nM- 500 nM) from midblastula (NF8.5) to late gastrula (NF12). qPCR analysis of RA target genes *hoxa1*.L (B), *hoxb1*.S (C), *cyp26a1*.L (D), and *aldh1l1* (E). Expression levels were normalized to control samples. Box-and-whisker (5-95 percentile) plots; statistical significance was determined using one-way ANOVA compared to the control expression.

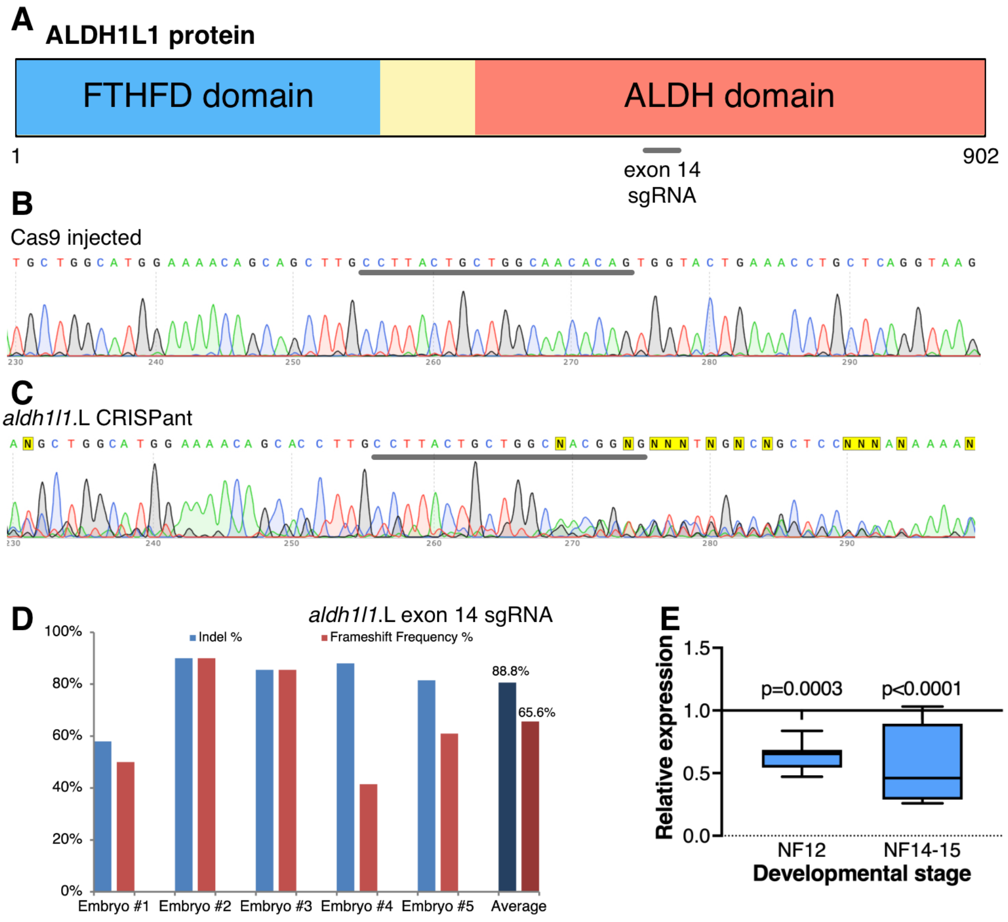


**Supplementary Figure 4. CRISPR/Cas9 targeting of the *aldh1l1* gene in *Xenopus* embryos.** (A) Schematic representation of the ALDH1L1 protein and its constituent domains. The position of the sgRNA used is shown. (B, C) Sequencing traces of genomic DNA extracted from control, Cas9-injected (B), and *aldh1l1* CRISPant (C) embryos at NF19-20. The sgRNA sequence is shown (line). (D) TIDE analysis for indels and frameshift mutations in the *aldh1l1* gene in five individual CRISPant embryos. (E) Expression change (qPCR) of the *aldh1l1*.L gene in CRISPant embryos during late gastrula (NF12) and early neurula (NF14-15). Box-and-whisker (5-95 percentile) plot; statistical significance was calculated using one-way ANOVA compared to the control expression.

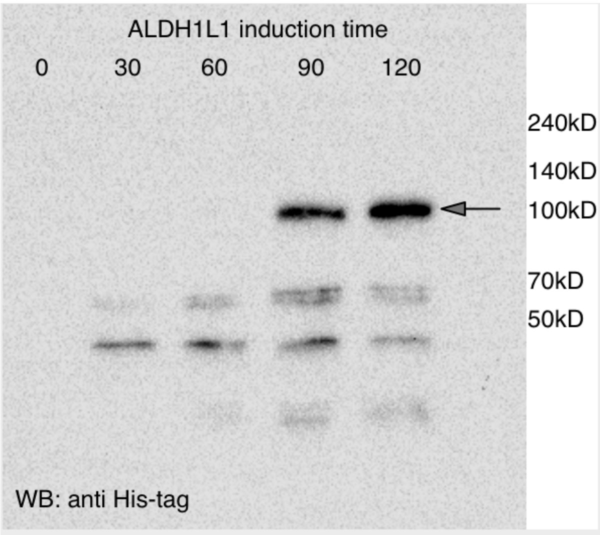


**Supplementary Figure 5. Production of the human ALDH1L1 enzyme in bacteria.** A cDNA, encoding the human ALDH1L1 enzyme was subcloned into the pET28a vector to produce the His-tagged enzyme in bacteria. Western blot analysis of the production of ALDH1L1 in bacteria as a function of time following induction with isopropyl β-D-1-thiogalactopyranoside (IPTG). The protein was detected with anti-His antibodies.

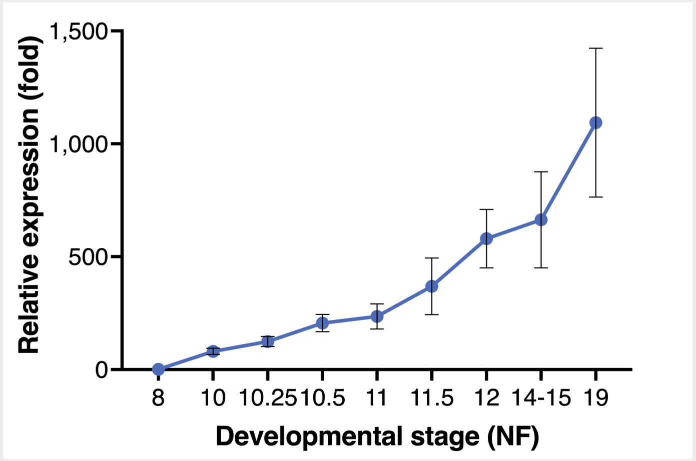


**Supplementary Figure 6. Temporal pattern of *aldh1l1* expression.** Control embryos were collected from blastula (NF8) to NTC stage (NF19). The temporal pattern of the *aldh1l1* gene expression was determined by qPCR of RNA extracted from these samples and analyzed across six embryo batches.

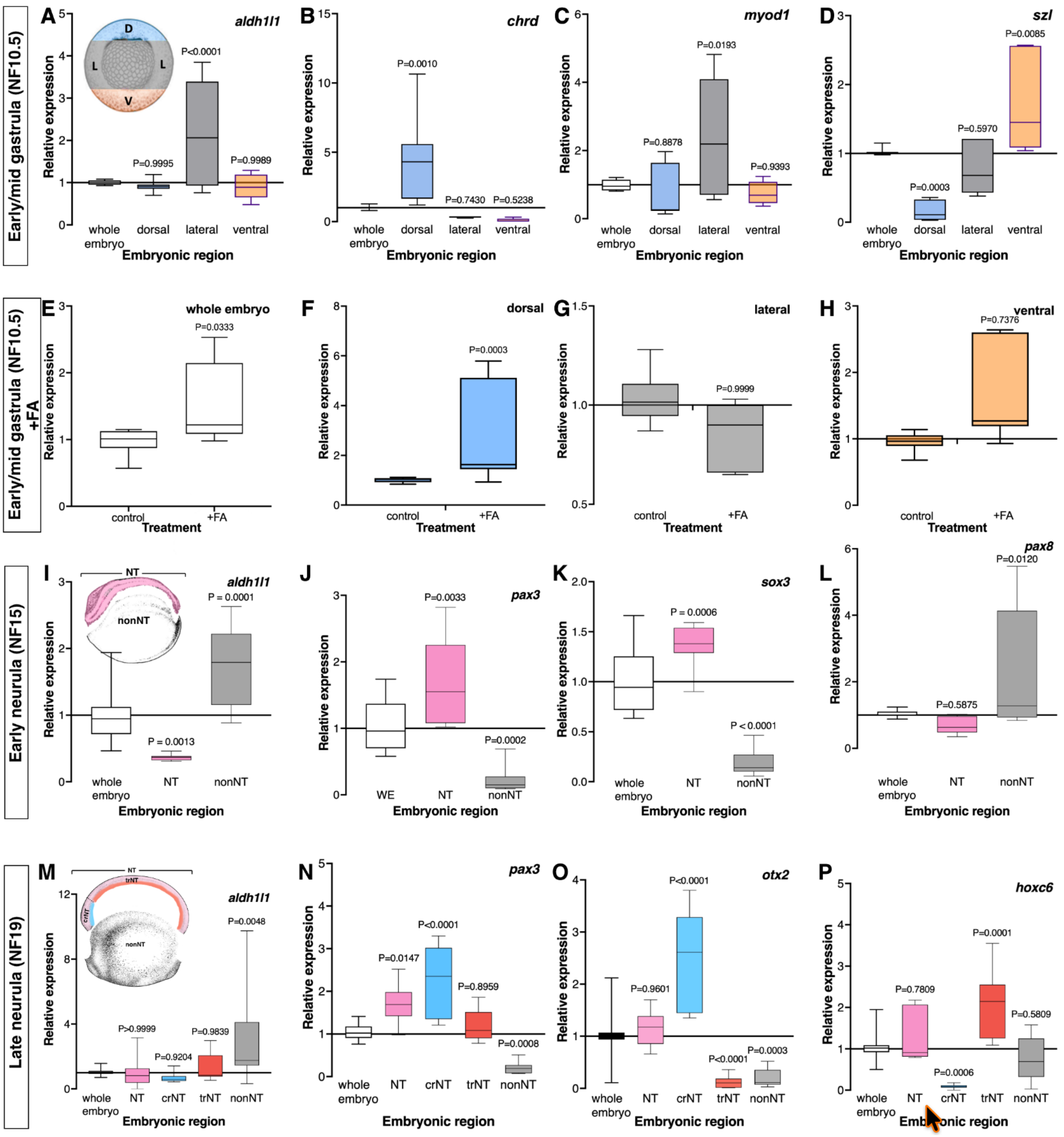


**Supplementary Figure 7. FA promotes the up-regulation of *aldh1l1* in the dorsal embryonic region.** (A-D) To determine the spatial localization of the *aldh1l1* transcripts during early/mid gastrula (NF10.5), embryos were dissected into dorsal, lateral, and ventral marginal zone regions, and RNA was prepared from each region. (A) qPCR analysis to determine the relative abundance of the *aldh1l1* transcripts in each region. Inset: schematic representation of the dissected regions. (B-D) Analysis of *chrd* (B), *myod1* (C), and *szl* (D) transcripts as dorsal, lateral, and ventral markers, respectively. (E-G) Dissected marginal zones were also prepared from FA-treated embryos and compared to the marginal zone fragments from control embryos. (E) Expression of *aldh1l1* in whole embryos. (F) Up-regulation of *aldh1l1* expression in DMZs. (G) Analysis of *aldh1l1* transcripts in LMZs. (H) Analysis of *aldh1l1* transcripts in VMZs. (I-L) Spatial distribution of *aldh1l1* transcripts during early neurula stages (NF15). Embryos were dissected into the prospective dorsal neural tube region (NT) and non-neural region (nonNT). (I) qPCR analysis of the *aldh1l1* transcripts in NT and nonNT regions. Inset: Schematic representation of the dissected regions. (J-L) Analysis of *pax3* (J), *sox3* (K) as neural markers, and *pax8* (L), as a lateral mesoderm marker. (M-P) Embryos were dissected during late neurula stages (NF19) to separate the dorsal neural tube region (NT) from the non-neural region (nonNT). Also, NT fragments were further dissected into cranial (crNT) and trunk (trNT) NT regions. (M) Expression of *aldh1l1* in the dissected regions. Inset: Schematic representation of the dissected regions. (N-P) Analysis of *pax3* (N), *otx2* (O), and *hoxc6* (P), neural, cranial neural, and trunk neural markers, respectively. Box-and-whisker (5-95 percentile) plots; statistical significance was calculated using one-way ANOVA.
